# Functional organization of motion and disparity sensitivity in human visual cortex

**DOI:** 10.1101/250878

**Authors:** Peter J. Kohler, Wesley Meredith, Anthony M. Norcia

## Abstract

Vision with two eyes makes perception of weak visual contrast signals easier and, due to the lateral separation of the eyes, allows for the triangulation of depth relationships. While binocular summation of contrast signals affords the observer increased sensitivity, binocular summation of spatial cues related to changes in depth is associated with decreased sensitivity to the corresponding retinal image displacements. Perceptual models of contrast and motion-in-depth sensitivity have explained this divergence in sensitivity by proposing that probabilistic neural noise limits summing and differencing operations on small signals. Because these models do not scale well for highly suprathreshold visual signals typical of the natural environment, we approached the question of how dynamic binocular image differences are coded using direct neural measurements. Here we use Steady-State Visual Evoked Potentials in human participants to show that inter-ocular differences in retinal image motion that produce elevated perceptual thresholds generate strongly suppressed evoked response amplitudes compared to motion that is matched between the two eyes. This suppression is strongly dependent on the availability of well-defined spatial references in the image and is highly immature in 5-month-old infants. Because the suppression is of equal strength for horizontal and vertical directions of motion, it is not specific to the perception of motion in depth. Relational image cues play a critical role in early to intermediate perceptual processing stages, and these results suggest that a succession of spatial and inter-ocular differencing operations condition the visual signal representation, prior to the extraction of motion-in-depth.

**Significance Statement:** The present work underscores the importance of relational spatial cues in both the motion and disparity domains for binocular visual coding. Relative motion and relative disparity cues not only support fine-grain displacement sensitivity but strongly influence suprathreshold responsiveness. Extraction of these cues supports a powerful binocular interaction within the motion pathway that is suppressive in nature and poorly developed in infants. This suppressive interaction is present for both horizontal and vertical directions of motion and is thus not specific to motion-in-depth, as previously believed, but is rather hypothesized to be a pre-processing step, with motion-in-depth being computed at a later or separate stage.

## Introduction

There is robust sensitivity to both direction of motion and retinal disparity in primary and higher-order visual cortex of primates. Direction tuning is present within the classical receptive field (Hubel and Wiesel, 1968; Dubner and Zeki, 1971; Dow, 1974; Schiller et al., 1976; Maunsell and Van Essen, 1983; Mikami et al., 1986; Orban et al., 1986), but can be modified by motion in the surround. These surround effects (Bridgeman, 1972; Allman et al., 1985; Born, 2000; Cao and Schiller, 2003; Shen et al., 2007) convey sensitivity to motion-defined discontinuities and to relative motion (where two or more velocities can be compared). Psychophysically, human observers are much more sensitive to relative motion than to absolute (unreferenced) motion (Legge and Campbell, 1981; Nakayama and Tyler, 1981; McKee et al., 1990).

Disparity tuning is also strongly present in V1 classical receptive fields (Poggio et al., 1985; Poggio et al., 1988; Cumming and Parker, 1997). Unlike the case just described for motion, the disparity of stimuli in the non-classical surround has little or no effect on V1 disparity tuning (Cumming and Parker, 1999; Bakin et al., 2000; Cumming and Parker, 2000; Samonds et al., 2017). By contrast, in V2 and beyond, responses to disparate stimuli within the classical receptive do depend on the disparities present in the surround, to varying degree (Thomas et al., 2002; Umeda et al., 2007; Anzai et al., 2011; Shiozaki et al., 2012). As for motion, psychophysical measurements indicate that human observers are much more sensitive to relative disparity than absolute disparity (Glennerster et al., 2002; Petrov and Glennerster, 2004, 2006).

Motion responses in V1 and V2 and other visual areas such as MT have been studied primarily for the case of stimuli that move in the fronto-parallel plane – somewhat of a special case, since objects move in three dimensions. With natural stimuli, there are two main cues that can be decoded to signal three-dimensional motion-in-depth (MID). The visual system can read out the binocular disparity of an object relative to the fixation plane and track how this information varies over time (change of disparity over time or CDOT). Another possibility is to compare object velocity from each monocular image (inter-ocular velocity difference or IOVD). Both cues provide partial, but not complete information for motion-in-depth (Lages and Heron, 2010).

Psychophysical work suggests that both IOVD and CDOT cues support percepts of motion in depth (Harris et al., 2008; Czuba et al., 2010; Cormack et al., 2017), but these experiments have been done in the context of scenes that do not have strong discontinuities in either motion or disparity. This approach may be limiting because of the importance of references for the precision of both motion and disparity discrimination, indicating that the visual system is highly adapted to detecting relative rather than absolute stimulus attributes.

Here we systematically explore the importance of clearly defined references on neural responses to motion and disparity using minimally complex scene structures comprised of regions defined by motion, disparity or both. We also take advantage of the fundamental asymmetry in retinal stimulation caused by the lateral separations of the eyes and compare stimuli with horizontally displaced motion signals that are ecologically relevant to MID, to stimuli with vertically displaced motion signals that are not. This allows us to separate neural responses specifically adapted to 3D-motion from those that support more generic image-processing functions. We find strong evidence for an effect of references on both motion and disparity responses and show that these reference-dependent responses are highly immature in 5-month-old infants. We also show strong equivalences between 3D-motion compatible and incompatible IOVD- and CDOT-based responses that logically arise before or in parallel with the perceptually relevant activity specific to 3D-motion.

## Results

Figure 1 shows the main stimulus configurations in schematic form. In each experimental condition, apparent motion at 2 Hz was presented in alternate bands of the display, with the amplitude of the displacement spanning 0.5 to 16 arcmin in 10 equal log steps for the adults (2-32 arcmin for infants). The adjacent, non-moving bands manipulated the availability of reference cues for motion, disparity or both. The different stimulus conditions manipulated whether the motion was in-phase or anti-phase between the two eyes, the availability and nature of motion and/or disparity cues in the non-moving reference bands, or the inter-ocular correlation of the moving bands. By making these comparisons systematically, we were able to evaluate how relative motion and disparity cues, inter-ocular phase and the CDOT and IOVD cues determine the properties of the displacement response function.

**Figure 1.**
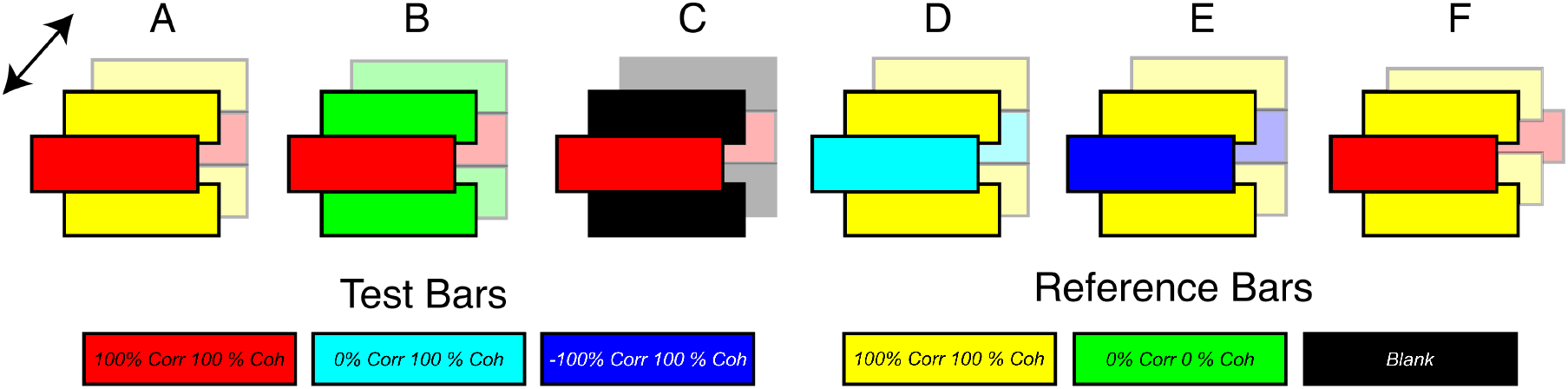
Schematic illustrations of the stimulus configurations. The displays comprised random dot kinematograms. Alternate bands (10 in the actual display) presented coherently moving dots whose motion was either in-phase between the two eyes or in anti-phase. The other set of 10 bands comprised different reference conditions. The moving bands are depicted here as a central band with the reference bands flanking it. The inter-ocular correlation in the moving bands was either 100% (red), 0% (cyan) or −100% (blue). The reference bands were either static and 100% correlated between eyes (yellow), temporally uncorrelated and inter-ocularly uncorrelated (green) or blank (black). Both horizontal and vertical directions of motion were presented. Note that in Experiment 3, the endpoints of the monocular apparent motion trajectories were manipulated such that moving bands alternated between equal and opposite values of crossed and uncrossed disparity (F)for the horizontal anti-phase condition and between right-hyper and right-hypo disparity for the vertical anti-phase condition.

### Absolute and relative motion responses

The Steady-State Visual Evoked Potential (SSVEP) related to both absolute (unreferenced) and relative (referenced) in-phase motion occurs at even harmonics of the stimulus frequency, with the second harmonic response (2F/4 Hz) being the largest (see Figure 2 for *2F* and Supplementary Figure 1 for *4F*/8Hz response functions).

**Figure 2.**
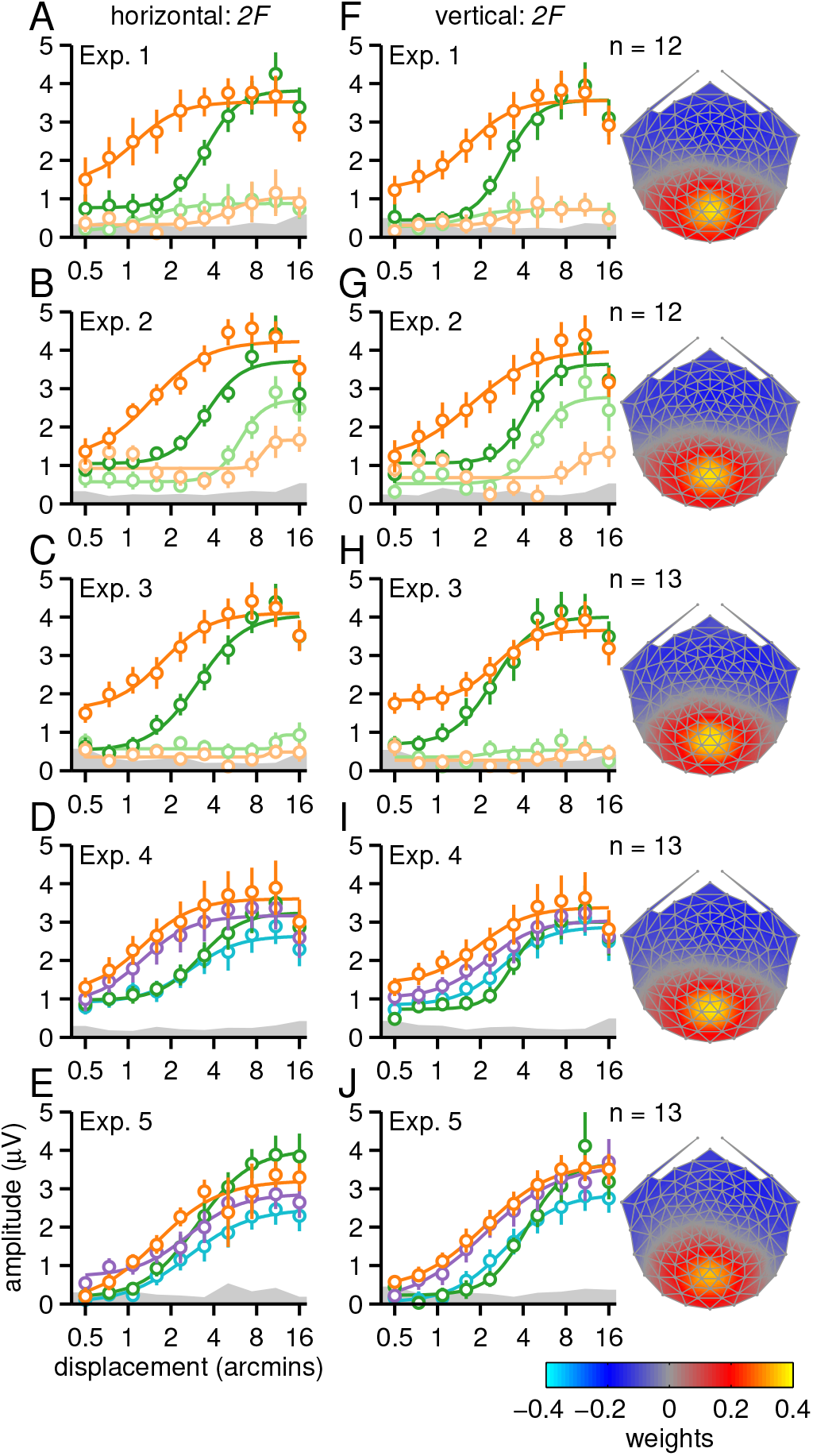
Second-harmonic response functions. Panels A-E depict displacement response functions for the horizontal direction of motion and panels F-J the vertical direction of motion. Referenced and unreferenced in-phase motion are shown in dark and light orange, while anti-phase motion is shown in dark and light green. Note that for Experiments 4 and 5, unreferenced conditions were replaced with uncorrelated and anti-correlated motion, shown in purple for in-phase and light blue for anti-phase. Smooth curves are Naka-Rushton function fits to the data (average fit parameters are shown in Supplementary Figure 2). The gray bands at the bottom of the plots indicate the background EEG noise level, with the top of the band indicating the average noise level across two neighboring side-bands, averaged across conditions. Error bars plot +/- 1 standard error of the mean (SEM). Data are from the first reliable component from an RC analysis run on 2F data from all conditions, separately for each experiment. Rightmost column shows the scalp topography of this component, which is centered over early visual cortex. Topographies were derived separately for each experiment, but are quite similar. See text for additional details.**Second-harmonic response functions**. Panels A-E depict displacement response functions for the horizontal direction of motion and panels F-J the vertical direction of motion. Referenced and unreferenced in-phase motion are shown in dark and light orange, while anti-phase motion is shown in dark and light green. Note that for Experiments 4 and 5, unreferenced conditions were replaced with uncorrelated and anti-correlated motion, shown in purple for in-phase and light blue for anti-phase. Smooth curves are Naka-Rushton function fits to the data (average fit parameters are shown in Supplementary Figure 2). The gray bands at the bottom of the plots indicate the background EEG noise level, with the top of the band indicating the average noise level across two neighboring side-bands, averaged across conditions. Error bars plot +/- 1 standard error of the mean (SEM). Data are from the first reliable component from an RC analysis run on 2F data from all conditions, separately for each experiment. Rightmost column shows the scalp topography of this component, which is centered over early visual cortex. Topographies were derived separately for each experiment, but are quite similar. See text for additional details.**Second-harmonic response functions**. Panels A-E depict displacement response functions for the horizontal direction of motion and panels F-J the vertical direction of motion. Referenced and unreferenced in-phase motion are shown in dark and light orange, while anti-phase motion is shown in dark and light green. Note that for Experiments 4 and 5, unreferenced conditions were replaced with uncorrelated and anti-correlated motion, shown in purple for in-phase and light blue for anti-phase. Smooth curves are Naka-Rushton function fits to the data (average fit parameters are shown in Supplementary Figure 2). The gray bands at the bottom of the plots indicate the background EEG noise level, with the top of the band indicating the average noise level across two neighboring side-bands, averaged across conditions. Error bars plot +/- 1 standard error of the mean (SEM). Data are from the first reliable component from an RC analysis run on 2F data from all conditions, separately for each experiment. Rightmost column shows the scalp topography of this component, which is centered over early visual cortex. Topographies were derived separately for each experiment, but are quite similar. See text for additional details.**Second-harmonic response functions**. Panels A-E depict displacement response functions for the horizontal direction of motion and panels F-J the vertical direction of motion. Referenced and unreferenced in-phase motion are shown in dark and light orange, while anti-phase motion is shown in dark and light green. Note that for Experiments 4 and 5, unreferenced conditions were replaced with uncorrelated and anti-correlated motion, shown in purple for in-phase and light blue for anti-phase. Smooth curves are Naka-Rushton function fits to the data (average fit parameters are shown in Supplementary Figure 2). The gray bands at the bottom of the plots indicate the background EEG noise level, with the top of the band indicating the average noise level across two neighboring side-bands, averaged across conditions. Error bars plot +/- 1 standard error of the mean (SEM). Data are from the first reliable component from an RC analysis run on 2F data from all conditions, separately for each experiment. Rightmost column shows the scalp topography of this component, which is centered over early visual cortex. Topographies were derived separately for each experiment, but are quite similar. See text for additional details.

In the first condition of Experiment 1, the moving test bands are flanked by adjacent reference bands containing static dots (Figure 1A) and the SSVEP amplitude is a saturating function of horizontal displacement (Figure 2A, dark orange). This response function is well-described by the Naka-Rushton function (Naka and Rushton, 1966) and fits of this function to the data are plotted as smooth curves in Figure 2 and elsewhere. In the second condition of Experiment 1, the static reference dots were replaced with dots that were temporally incoherent (Figure 1B), making it impossible to calculate a unique relative velocity because the reference bands contain a very broad and random distribution of velocities. This manipulation strongly reduces the response amplitude for in-phase motion (Figure 2A, light orange). In the vertical direction, in-phase motion produced very similar *2F* responses, with similar differences between referenced and unreferenced conditions (Figure 2F).

The incoherent dots in the reference bands in Experiment 1 may have masked the moving dots through suppressive lateral interactions. We tested for this in Experiment 2 by recording from a second group of participants using a display in which the moving bands were the same as in Experiment 1 but the reference bands were blank (Figure 1C). We also included the referenced motion conditions of Experiment 1 for a within-observer comparison. We found that the unreferenced response was now measurable, but a large difference in amplitude persisted between referenced and unreferenced in-phase motion for both horizontal (Figure 2B) and vertical conditions (Figure 2G).

Experiment 3 used the same conditions as Experiment 1, but the endpoints of the monocular apparent motion trajectories were manipulated (see Figure 1F). This has a strong effect on the percept and first harmonic responses produced by the anti-phase conditions (see below), but produces *2F* responses that are quite similar to Experiment 1, for all conditions. We again saw much weaker responses for unreferenced compared to referenced in-phase motion for both horizontal and vertical conditions (see Figure 2C, H).

We tested the significance of the difference between referenced and unreferenced in-phase motion with jackknifed t-tests on the Naka-Rushton function fit parameters (see Supplementary Figure 2 for a summary of the fit parameters and Table S1 for results). For the horizontal conditions, the *R_max_* parameter was significantly larger for referenced motion in Experiment 2, while the baseline *(b)* parameter was significantly larger in Experiments 1 and 3. In addition, there was a marginally significant trend towards higher sensitivity (lower *d_50_*) for the referenced condition in Experiment 2 (*p* = 0.058). For the vertical conditions, the *R_max_* parameter was significantly larger for Experiments 1 and 3, and there was a similar trend in Experiment 2 (*p* = 0.053), while the baseline parameter was significant in Experiment 3 and marginally significant in Experiment 1 (*p* = 0.052). The larger *R_max_* and baseline parameters, and in one case lower *d_50_*, for the referenced compared to the unreferenced conditions (see Table S1), demonstrate that our paradigm is sensitive to relative-motion-specific responses.

### Sensitivity to inter-ocular phase

By making the inter-ocular phase the only difference between conditions, we can focus the comparison of responses on specifically binocular mechanisms. Anti-phase horizontal motion in the two eyes creates IOVD and CDOT cues that are not present in the in-phase conditions. When the motion direction is horizontal, these cues support a percept of motion-in-depth.

In the presence of a static reference, anti-phase motion evoked a response that is a saturating function of displacement. As for in-phase motion, unreferenced responses were strongly reduced (compare dark and light green in Figure 2A, F). We again tested for differences between referenced and unreferenced conditions using jackknifed *t*-tests on the fit parameters (see Table S2 for results). For the horizontal conditions, the reference effect manifested as larger *R_max_* in Experiments 1 and 3, and as larger baselines in Experiment 2. In addition, *d_50_* was lower for the referenced condition in Experiments 2 and 3, but higher in Experiment 1. For the vertical conditions, the differences manifested as larger *R_max_* for the referenced condition in Experiments 1 and 3, and as larger baselines in Experiment 2.

The response functions for the referenced anti-phase conditions were shifted rightward on the displacement axis relative to the in-phase conditions, suggesting that the visual system was less sensitive to anti-phase motion in the two eyes. We assessed the significance of the difference between the in-phase and anti-phase conditions with jackknifed *t*-tests on the fit parameters (see Table S3 for results). Note that the five adult experiments (1, 2, 3, 4 and 5) all had virtually identical referenced in-phase and antiphase conditions, so we compared the fit parameters for all of them. The *d_50_* parameter was significantly lower for in-phase compared to anti-phase motion for the horizontal conditions in all five experiments, and for the vertical conditions in four out of five, with Experiment 3 as the exception. The *R_max_* parameter was only significant in Experiment 3, were *R_max_* was smaller for in-phase than anti-phase for both horizontal and vertical conditions. The exponent (*n*) parameter was smaller for in-phase than anti-phase in the vertical conditions of Experiments 1, 2 and 5 and there was a similar trend in Experiment 4 (*p* = 0.065). There were no significant exponent effects for the horizontal conditions, although Experiment 2 approached significance (*p* = 0.084). The baseline parameter was larger for in-phase in the horizontal conditions in Experiment 3 and in the vertical conditions in Experiments 1, 3 and 4. The differences between in-phase and anti-phase responses are thus relatively stable across the referenced conditions that were repeated in multiple experiments.

In Experiment 2, where the static dot reference is replaced with a blank reference, the anti-phase response function is no longer shifted to the right of the in-phase response function, but is rather shifted to the left (Figure 2B, G). This manifests as a larger *R_max_* for the horizontal conditions, with a trend towards significance for the vertical conditions (*p* = 0.10; see Table S4). This result, when compared to the results of the other experiments, indicates that the suppression of responses to anti-phase motion, relative to responses to in-phase motion, depends on the content of the reference region.

### Relationship to perceptual stereo-movement suppression

We have previously observed a reduced amplitude evoked response to anti-phase compared to in-phase motion (Cottereau et al., 2014). We suggested that this effect may be related to the perceptual phenomenon known as “stereo-movement suppression” (Tyler, 1971), a reduction in displacement sensitivity under binocular viewing conditions that has been replicated numerous times and is usually attributed to the stereoscopic motion system (Tyler, 1971; Beverley and Regan, 1973; Brooks and Stone, 2006; Katz et al., 2015; Cooper et al., 2016).

In our measurements, referenced anti-phase responses are reduced relative to inphase responses both for horizontal conditions, which elicit stereoscopic motion, and vertical conditions, which do not. To determine whether our displays also elicit perceptual suppression, we conducted two psychophysical motion detection experiments, using the method of ascending and descending limits. In the first, participants viewed the static reference conditions from Experiments 1 and 2. In the second, participants viewed the blank reference conditions from Experiment 2. To directly link perceptual data to the SSVEP response functions, we recorded SSVEPs during the psychophysical measurements. The SSVEP data from both psychophysical experiments were projected through the first reliable component generated by the RCA done on the *2F* data from Experiment 2, and averaged over ascending trials and flipped versions of the descending trials.

In the first psychophysical experiment, the range of displacements was decreased to 0.25 to 4 arcmin to place the start of the ascending sweep (and the end of the descending sweep) below perceptual threshold (see Figure 4). The SSVEP response functions resembled those of previous experiments. Response amplitudes were higher for in-phase than for anti-phase motion (compare dark green and orange in Figure 2 with Figure 4A, B). This manifested as lower *d_50_* for in-phase compared to anti-phase (0.83 vs 1.58 for horizontal, 0.83 vs 1.75 for vertical) and higher *R_max_* (1.50 vs 0.60 for horizontal, 1.35 vs 0.26 for vertical), although significant differences could not be obtained consistently due to noisy fits of the near-baseline anti-phase responses. Psychophysical thresholds, averaged over ascending and descending sweeps, were higher for anti-phase motion by factors ∼1.5 for horizontal and 2 for vertical displacements (Figure 4C), indicating that our experimental conditions do give rise to the stereo-movement suppression phenomenon. Paired *t*-tests on individual participant thresholds confirmed that these differences were significant for both horizontal (*t*(11) = −4.40, *p* = 0.001) and vertical (*t*(11) = −7.93, *p* < 0.0005) displays.

The second psychophysical experiment used the same range of displacements as the main experiments, under the assumption that unreferenced thresholds would be higher. This is in fact what we observed: SSVEP thresholds across conditions, captured by the *d_50_* parameter, were on average higher for the unreferenced data by a factor of ∼5.3, compared to the referenced data from the first psychophysical experiment. The SSVEPs replicated the reversal observed in the results from Experiment 2 (compare Figure 2B, G with Figure 4D, E). The *R_max_* value was significantly lower for in-phase compared to anti-phase in the vertical conditions (1.38 vs 2.79; *t*(10) = −2.70, *p* = 0.022) and there was a similar trend in the horizontal conditions (1.58 vs 2.28; *t*(10) = −2.04, *p* = 0.069). There was also a trend towards lower *d_50_* for anti-phase compared to in-phase in the vertical conditions (6.38 vs 4.58; *t*(10) = 2.04, *p* = 0.069), but no evidence of this for the horizontal conditions (5.69 vs 5.76; *t*(10) = −0.10, *p* = 0.92). In addition, the exponent was significantly larger for in-phase compared to anti-phase (*t*(10) = 2.52, *p* = 0.030). The psychophysical thresholds for the unreferenced data, averaged over ascending and descending sweeps, were higher than for the referenced data by a factor of ∼4, but led to comparable differences between in-phase and anti-phase motion. Anti-phase thresholds were higher than in-phase by factors ∼1.75 for horizontal and ∼1.5 for vertical (Figure 4C), indicating that the perceptual stereo-movement suppression phenomenon persisted in the blank reference condition. Paired *t*-tests on individual participant thresholds confirmed that these differences were significant for both horizontal (*t*(10) = −7.03, *p* < 0.0005) and vertical (*t*(10) = −5.36,*p* < 0.0005).

These results of the first psychophysical experiment extend the pattern seen over the larger range of displacement amplitudes in Figure 2 to the threshold regime. For both horizontal and vertical directions of motion, SSVEPS are weaker and perceptual thresholds are higher for anti-phase compared to in-phase conditions. Because the vertical conditions do not give rise to a percept of motion-in-depth, we conclude that the suppressed anti-phase responses are not uniquely associated with percepts of motion-in-depth. In the second psychophysical experiment, the SSVEP data replicate the results of Experiment 2, with overall weaker responses for unreferenced conditions, and slightly stronger responses for anti-phase than in-phase, for both horizontal and vertical conditions. Psychophysical thresholds are higher for unreferenced than referenced conditions, but in contrast to the SSVEP data, thresholds are higher for anti-phase conditions than in-phase. This discrepancy can perhaps be explained if participants relied on subtle reference cues in the experimental environment, that were not encoded by most of the neurons generating the population response measured by the SSVEPs. Nonetheless, the SSVEP data are consistent with the conclusion of Experiment 2: the suppression of response seen with anti-phase motion depends on the content of the reference region and is independent of whether the displays are horizontal or vertical.

### Binocular immaturity in infants

When we presented the horizontal referenced and unreferenced in-phase motion displays to 18 infants (∼5 months old), we found that their *2F* responses were independent of whether the reference bands contained static dots or were blank (see Figure 4). *t*-tests comparing the fit parameters for the referenced and unreferenced conditions were far from significance for both in-phase and anti-phase conditions (*p*’s for all parameters > 0.2). For the referenced conditions, the in-phase stimulus had a lower *d_50_* than the antiphase stimulus (*t*(17) = 2.55, *p* = 0.021), while in the unreferenced conditions, the baseline was lower for in-phase than anti-phase, although not quite significantly so (*t*(17) = −1.90, *p* = 0.074).

There are several differences between the infant and adult data: First, infant *2F* responses are not increased in the presence of a reference, in distinct contrast to the adult data (see Figure 2). Second, infant anti-phase *2F* responses are larger than in-phase responses, in a reversal of the adult response pattern. Note that we observed similar reversals for adults in Experiment 2 when the reference was blank (see Figure 2B, G) and in the second psychophysical experiment (see Figure 3). The infant response pattern with the full-cue, referenced display thus resembles that of the adults in the blank reference conditions. In both the in-phase and anti-phase motion conditions, the infant response shows no measureable effect of the reference. Moreover, there is no evidence for suppression of referenced anti-phase responses relative to in-phase ones, indicating infants have a specifically binocular immaturity in sensing inter-ocular phase.

**Figure 3.**
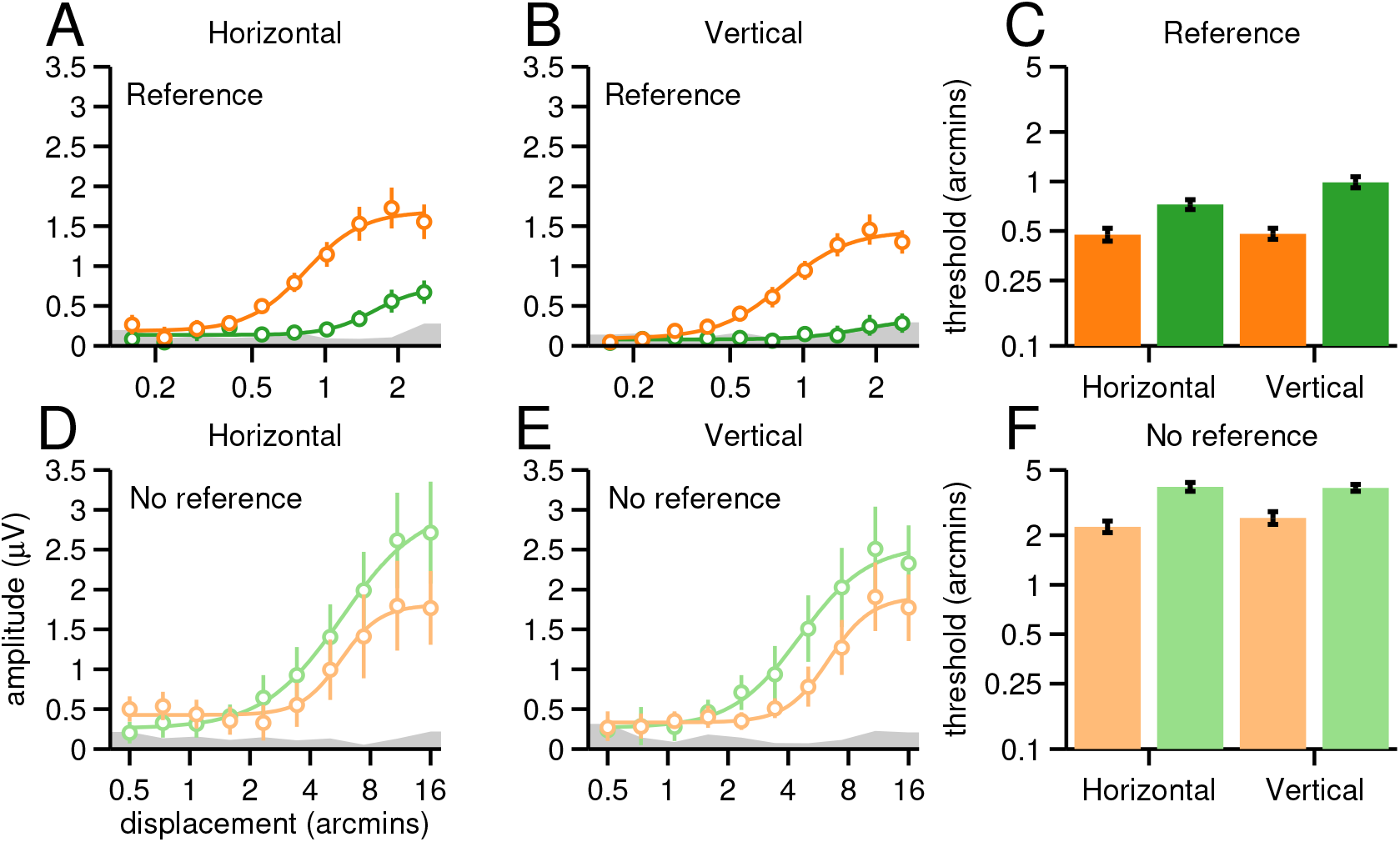
SSVEP response functions and psychophysical detection thresholds. Panels A and B show SSVEP data from horizontal and vertical direction of motion trials with a static reference (n = 12). Inphase conditions are plotted in orange and anti-phase conditions in green. We ran both descending and ascending displacement sweeps; response functions are averages over ascending and flipped versions of descending trials. As in Figure 2, gray bands represent the average background EEG noise and smooth curves are Naka-Rushton fits to the data. Responses were weaker for anti-phase compared to in-phase for both horizontal and vertical motion. Panel C shows psychophysical thresholds for in-phase (orange) and anti-phase (green) conditions for horizontal and vertical directions of motion, plotted on a log scale, and again averaged over ascending and descending sweeps. Thresholds were higher for anti-phase compared to in-phase motion. Panels D and E show SSVEP data for blank reference conditions (n = 11). Note that a larger range of displacements was used for the blank reference conditions and that the response functions depart from the noise level at higher displacements than in the static reference conditions. Unlike the referenced conditions, responses are weaker for in-phase compared to anti-phase. Panel F shows psychophysical thresholds for the blank reference condition. Overall psychophysical thresholds are higher by a factor of ∼4 than for the referenced conditions and thresholds are higher for anti-phase than in-phase motion for both orientations.

### Anti-phase suppression is a property of the IOVD system

The perceptual stereo-movement suppression effect has been explored in the context of stimuli that have both IOVD and CDOT cues (Tyler, 1971; Brooks and Stone, 2004; Harris et al., 2008; Katz et al., 2015; Cooper et al., 2016). In the experiments just described, IOVD and CDOT are present in both horizontally and vertically oriented displays, but only the horizontal versions of the two cues can support a computation of motion-in-depth (Lages and Heron, 2010). Given this asymmetry, it is not surprising that the perceptual literatures on CDOT and IOVD have focused on the horizontal case. The neural signature of anti-phase suppression we see in our data can be measured for both orientations. Because IOVD is an explicitly motion-based cue, we wanted to determine whether IOVD cues alone could elicit the suppression effects in our paradigm.

We isolated the IOVD cue using two approaches developed previously for this purpose. In Experiment 4, we used dots in the moving regions that were uncorrelated between the two eyes (Maeda et al., 1999; Shioiri et al., 2000; Nefs and Harris, 2010). In Experiment 5, we used dots that were anti-correlated between the two eyes (Rokers et al., 2008, 2009). The suppression effects were weaker and less consistent for the IOVD isolating conditions, but we nonetheless saw evidence that anti-phase suppression can be generated by the IOVD cue alone (see Table S5). For the uncorrelated horizontal conditions (purple and blue in Figure 2D), the anti-phase suppression manifested as lower *d_50_*, as well as a trend towards larger *R_max_ (p* = 0.080). A similar, but attenuated, pattern was seen for the uncorrelated vertical conditions (Figure 2I), where only the *d_50_* parameter came close to significance (*p* = 0.124). For the anti-correlated horizontal conditions (Figure 2E), the anti-phase suppression manifested as higher baseline for inphase, while for the vertical conditions there were trends towards lower *d_50_ (p* = 0.086) and higher *R_max_ (p* = 0.119) for in-phase.

Weaker suppression effects may occur because the uncorrelated and anti-correlated conditions cause a different mixture of disparity- and motion-related responses. The anti-correlated cue is expected to activate disparity-tuned cells in early visual cortex, with an inverted sign (Cumming and Parker, 1997). This would not be expected for the uncorrelated case. The two conditions may thus cause a different mixture of disparity- and motion-related responses. Moreover, both IOVD isolating conditions can trigger binocular rivalry that may reduce the response magnitude in the in-phase condition that is used as the reference for the suppression effect (compare orange and purple in Figure 2D, E, I and J). Nonetheless, the overall effects and trends in the data indicate that anti-phase suppression can be generated by the IOVD cue alone.

### Candidate signal related to MID from IOVD

The evoked response is comprised of multiple even and odd-harmonic response components and thus far we have concentrated on the 2^nd^ harmonic component. The 2^nd^ harmonic behavior was similar for horizontal and vertical directions of motion and it is thus not a likely source of MID signals for perception. To look further for a possible marker for evoked responses that could contribute to MID from IOVD we examined both the *2F* and *4F* SSVEPs for evidence of differential responses to horizontal and vertical directions of anti-phase motion under the assumption that response components that differed for perceptually relevant and irrelevant directions of motion could provide a substrate for MID from IOVD. We combined the data from the horizontal and vertical anti-phase conditions, recorded with the static reference, across Experiments 1, 2, 3 and 4, yielding a data set with 39 participants (note that only one session was added to the combined data for individuals that took part in several experiments). We then derived reliable components over the larger group separately for the *2F* and *4F* data, using the same RCA approach as for the individual experiments. Figure 5 shows *2F* (A) and *4F* (B) response functions for the two orientations of antiphase motion, from the first reliable component. The horizontal response functions are plotted in green, with the vertical data in red. The component topographies are plotted in (C) and (D). As expected, the *2F* topography is very similar to the *2F* topographies done separately for each experiment (compare Figure 2 and Figure 5C), while the *4F* topography was perhaps slightly broader (Figure 5D). There was no measurable difference between responses to horizontal and vertical orientations at *2F*, whereas for *4F* the green curve is shifted to the left, indicating that for the perceptually relevant horizontal stimulus orientation, responses rise out of the noise at lower displacements. This *4F* response difference manifested in lower *d_50_* for horizontal compared to vertical (*t*(38) = −2.71, *p* = 0.010). For *2F*, there was a trend towards a difference, but indicating higher sensitivity (lower *d_50_*) for vertical than horizontal (*t*(38) = 1.54, *p* = 0.132), and thus in the opposite direction expected of a MID-related signal. All other comparisons for both harmonics were far from significance (all *p*’s > 0.2).

**Figure 4.**
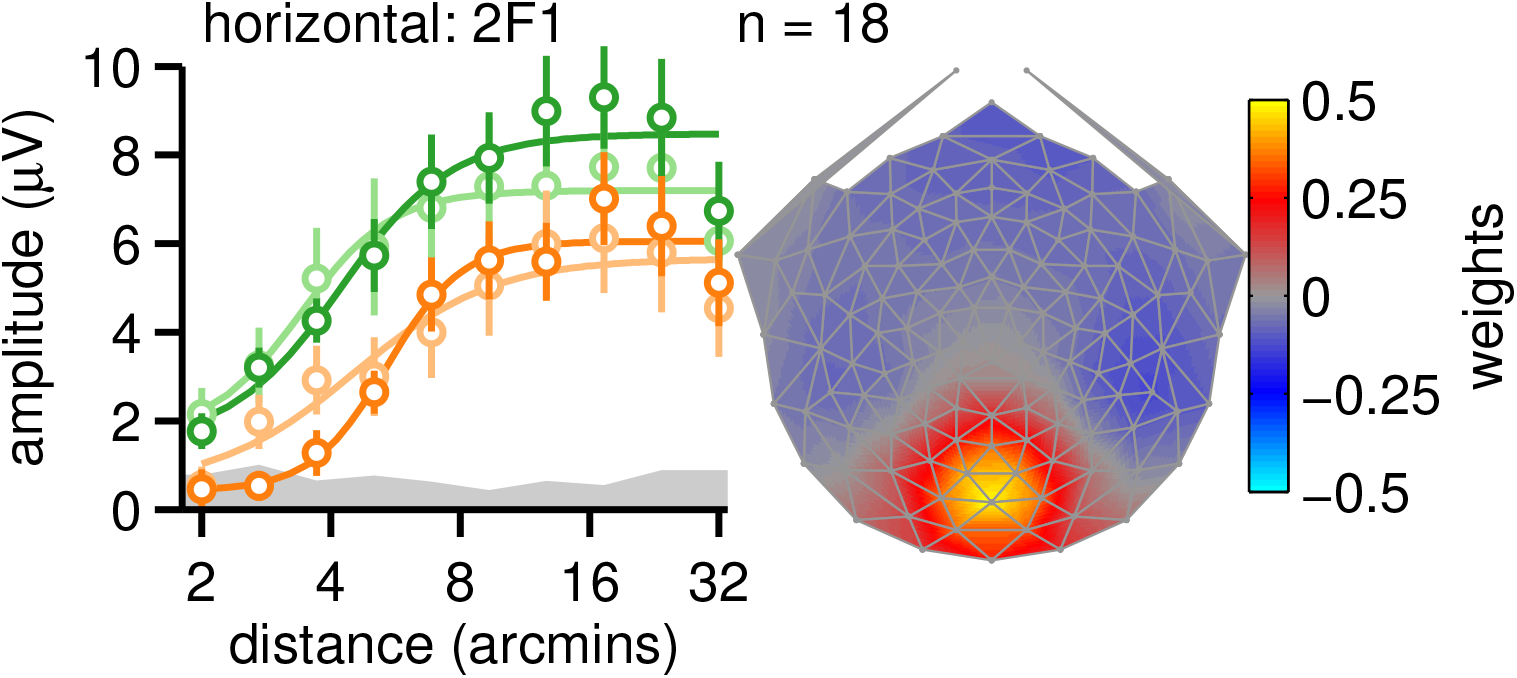
Infant second-harmonic response functions. In-phase motion response functions are shown in dark orange for the static reference condition and in light orange for the blank reference condition. Antiphase response functions are shown in dark green for the static reference condition and in light green for the blank reference condition. Smooth curves are Naka-Rushton fits to the data. Error bars are +/−1 SEM and the gray band indicates the average EEG noise level across conditions. Unlike adults, in-phase response functions lie to the right of anti-phase functions and do not show an effect of the reference. Right panel shows the scalp topography of the first (most) reliable component which is centered over early visual cortex.

**Figure 5.**
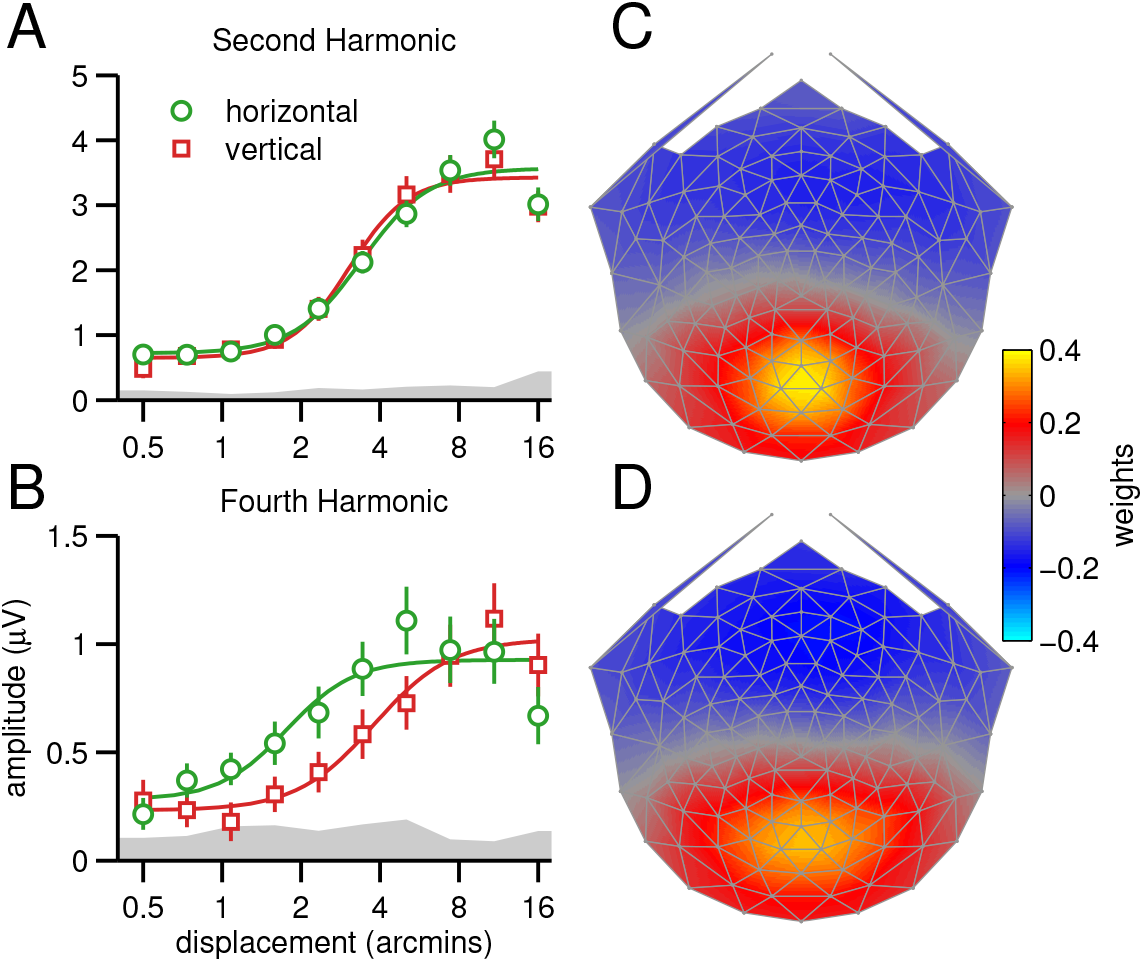
Candidate MID signal from IOVD. Response functions for horizontal (green) and vertical (red) anti-phase motion conditions, averaged across Experiments 1, 2, 3 and 4. The response functions are from the first reliable component of RCA done separately on 2F (A) and 4F (B) data, with the topographies shown on the right (C and D). Responses do not differ for the two directions of motion at 2F, but are different at 4F1 at the smaller displacements. An analogous analysis done on the uncorrelated and anti-correlated anti-phase conditions from Experiments 4 and 5 produced a similar pattern of results (see Supplementary Figure 3). Smooth curves are Naka-Rushton fits and gray bands are the averaged EEG noise level.

If the *4F* effect is in fact a substrate for MID from IOVD, we would expect it to replicate in the uncorrelated and anti-correlated IOVD-isolating conditions. To test this, we repeated the analysis with combined data from the horizontal and vertical anti-phase conditions from Experiments 4 and 5, yielding a data set with 22 participants. The first reliable components for *2F* and *4F* were similar to those produced for the larger data set (Supplementary Figure 3C and D), and the overall trend in the SSVEP response functions was similar (Supplementary Figure 3A and B), although the *d_50_* parameter was not quite significantly lower for horizontal than vertical (*t*(21) = −1.52, *p* = 0.144).

Overall, these results are consistent with the hypothesis that neurons generating the *4F* response can support a percept of motion in depth, but this would need to be tested by experiments demonstrating co-variation of the *4F* response with depth percepts.

### CDOT, IOVD, MID and image segmentation

The in-phase motion conditions consist of left/right or up/down motion in the plane, and are thus phenomenologically symmetric over time. Consistent with this, the evoked response is dominated by *2F* and higher even harmonics. By contrast, the anti-phase condition is phenomenologically asymmetric - for horizontal motion, the observers’ percept alternates between a segmented field of disparate bands and a field comprised of a flat plane (zero disparity over the whole image). This asymmetry in perceptual organization manifests in the evoked response as the presence of a response at the first harmonic (*1F*) of the stimulus frequency, i.e. the rate at which the perceptual organization changes (2 Hz). The *1F* displacement response functions are shown in Figure 6 for the five main experiments. The data are from the first reliable component produced by RCA performed separately on the *1F* data, using the same approach as for *2F*.

**Figure 6.**
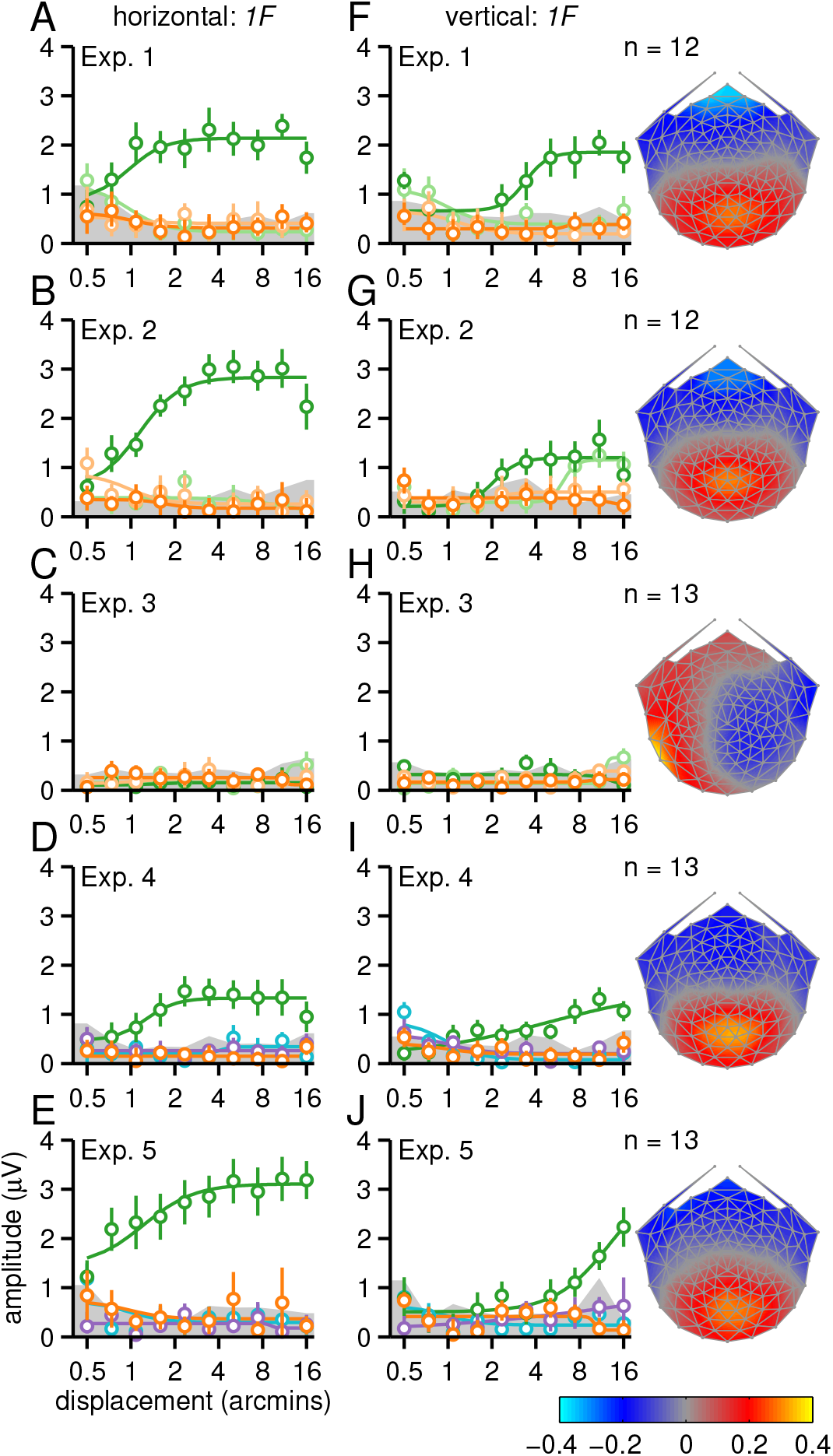
First-harmonic response functions. Panels A-E depict displacement response functions for the horizontal direction of motion and panels F-J the vertical direction of motion. Referenced and unreferenced in-phase motions are shown in dark and light orange, while anti-phase motion is shown in dark and light green. Note that for Experiments 4 and 5, unreferenced conditions were replaced with uncorrelated and anti-correlated motion, shown in purple for in-phase and light blue for anti-phase. Smooth curves are Naka-Rushton function fits to the data and gray bands at the bottom of the plots indicate the average background EEG noise level. Error bars plot +/− 1 standard error of the mean (SEM). Data are from the first reliable component from an RC analysis run on IF data from all conditions, separately for each experiment. The rightmost column shows the scalp topography of this component, which is centered over early visual cortex. Topographies were derived separately for each experiment but are quite similar, except for Experiment 3 where there is no reliable 1F response. See text for additional details.

Like the *2F* responses, the *1F* response is a saturating function of displacement, starting from the smallest displacement amplitude that depends on the presence of a local reference of static dots. In the horizontal referenced conditions of Experiments 1, 2, 4 and 5 there is strong *1F* response for anti-phase motion, (Figure 6A, B, D, E, dark green). This response is obliterated when the reference is replaced with uncorrelated dots or removed (light green), and in the in-phase conditions where there is no motion in depth (dark and light orange). Similar dependence on a correlated zero disparity reference has been observed with dynamic random dot displays that fully isolated the CDOT cue (Norcia et al., 2017a).

We also obtained measurable *1F* responses for vertical conditions with a static dot reference, but the response function is shifted to the right by a factor of ∼4 (Figure 6F, G, I and J, dark green). Here the stimulus contains vertical relative disparity. That this response is indeed a relative disparity response is indicated by the weak or absent *1F* responses in the uncorrelated and blank reference conditions (Figure 6F and G, light green). A vertical relative disparity signal has also been found with pure CDOT dynamic random dot stimuli (Norcia et al., 2017a). In both cases, displacement sensitivity is about 4 times better for horizontal than for vertical disparity.

The importance of references is well known for horizontal disparity (Westheimer and McKee, 1979; Glennerster and McKee, 1999; Petrov and Glennerster, 2006). The present results suggest that relative disparity is also calculated along the vertical direction, consistent with other recent findings (Norcia et al., 2017a), and provide support for a previous psychophysical finding that vertical disparities can be used for discontinuity detection (Serrano-Pedraza et al., 2010).

If *1F* is in fact due to the phenomenological asymmetry over time as the antiphase display alternates between uniform and segmented percepts, the first harmonic should disappear with a stimulus configuration in which the display no longer alternates between uniform and segmented states. Such a configuration also tests the alternative hypothesis that the asymmetry leading to *1F* is due to different response amplitudes for motion towards and away from the observer. For Experiment 3, we generated such a display by making the two endpoints of the monocular apparent motion trajectories straddle zero disparity, which means that the bands alternated between equal and opposite values of crossed and uncrossed disparity. This subtle manipulation eliminates the asymmetry associated with alternating between uniform and segmented percepts, but still gives rise to motion towards and away from the observer in the horizontal anti-phase conditions, and to left/right or up/down motion in the plane in the in-phase conditions. The latter point explains why *2F* responses are so similar between Experiments 1 and 3. The *1F* response is eliminated in Experiment 3, for both horizontal and vertical directions of motion, tying the response to asymmetric processing of the uniform vs segmented stimulus states (Figure 6C, H).

In the full-cue condition, the *1F* response could in principle arise from either the CDOT or IOVD cue, as both are present. However, the *1F* response becomes unmeasurable in the uncorrelated (Figure 6D, I) and anti-correlated (Figure 6E, J) conditions that eliminate the CDOT cue, indicating that *1F* responses are driven by CDOT. The fact that this CDOT-driven *1F* response can be measured for vertical relative disparities suggests that it is not exclusively a MID signal, at least at large disparity values.

Finally, in the case of the infants, a weak, but measurable *IF* response was present in the referenced horizontal condition (Figure 7, dark green). The sensitivity to displacement in our full cue display here is similar to previous measurements with CDOT-isolating dynamic random-dot stereograms (Norcia et al., 2017a). As for adults, the infant *1F* depends on the presence of the static dot reference and disappears when the blank reference is used (Figure 7, light green), although caution is needed here given the weak responses to the full-cue condition.

**Figure 7.**
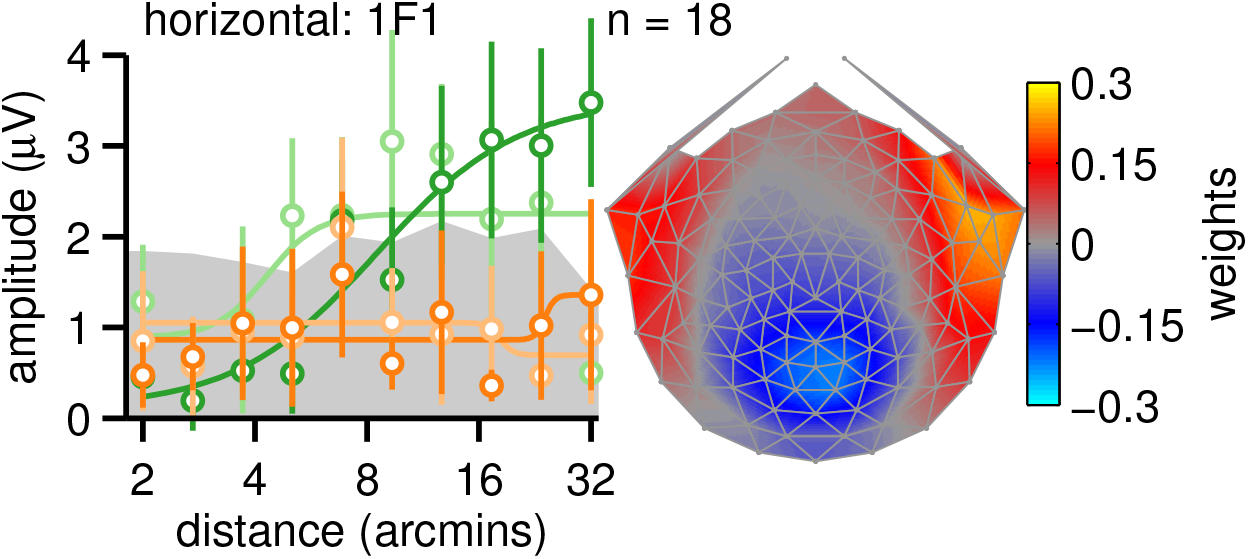
Infant IF SSVEP amplitude vs displacement response functions. In-phase motion response functions are shown in dark orange for static reference condition and in light orange for blank reference conditions. Anti-phase response functions are shown in dark green for the static reference condition and in tight green for the blank reference condition. Smooth curves are fits from a Naka-Rushton fimction. Error bars are +/−1 SEM, and the gray band indicates the average EEG noise level. The response topography is shown on the right. Unlike previous plots, data are plotted for the fifth reliable component because its topography was most similar to the topography of the adults. Note that the sign of the weights is arbitrary.

## Discussion

Our data provide new insights into the early stages of lateral motion and motion-in-depth processing. First, for the case of in-phase, lateral motion in the fronto-parallel plane, we show that both threshold and supra-threshold responsivity is strongly dependent on the presence of a nearby reference in adults, but not in 5-month-old infants. Infants were not sensitive to relative motion under the stimulus conditions we used. Single-unit recordings with large amplitude motions in cat lateral supra-sylvian cortex (von Grunau and Frost, 1983), pigeon tectum (Frost and Nakayama, 1983) and macaque areas MT, MST, superior colliculus and V1 (Bridgeman, 1972; Allman et al., 1985; Born, 2000; Cao and Schiller, 2003; Shen et al., 2007) have found cells that respond best when motion in the classical receptive field is surrounded by differential motion. The pattern of activity we observe is consistent with relative motion being processed via directionally opponent interactions between the classical receptive field and its non-classical surround. The lack of reference effects in infants in the present experiments may reflect an immaturity in these interactions.

Our results on anti-phase motion have identified an IOVD-based mechanism that is present for both horizontal and vertical directions of motion. The functional manifestations of this mechanism are an elevation of perceptual threshold for anti-phase motion relative to in-phase motion and a decrease in SSVEP response amplitude. Amplitude reductions were demonstrable with IOVD-isolating stimuli, linking the phenomenon to that cue. Because this suppression effect is equal for horizontal and vertical directions of motion, it is unlikely to be related to the extraction of MID, per se because stimulus information for MID from IOVD is only present for horizontal or near-horizontal directions of motion (Lages and Heron, 2010). Prior descriptions of the phenomenon as reflecting stereoscopic depth movement (Tyler, 1971) thus need to be revised. The observed suppression either precedes the computation of MID from this cue or operates in parallel.

In our anti-phase conditions, we presented opposite directions of motion to the two eyes and a differential response between in-phase and anti-phase conditions could thus only be generated after binocular combination. The anti-phase suppression we have observed is consistent with a dichoptic, directionally-opponent interaction. The functional form of the suppressive interaction is a rightward shift of the response curve on the displacement axis, consistent with a form of divisive normalization (Carandini, 2012). Prior work has suggested that the perceptual stereo-movement suppression effect is due to limits in sensitivity imposed by increased noise in binocular differencing operations (Brooks and Stone, 2004; Katz et al., 2015; Cooper et al., 2016). By using a direct neural measure over both threshold and supra-threshold levels, we see that the stimulus-driven population response itself is strongly attenuated. This attenuation is difficult to accommodate within a probabilistic, noise-limited detection framework. Suppression is more consistent with mutual inhibition between the two eyes, as initially suggested in the first observations of the suppression effect (Tyler, 1971).

Prior psychophysical work on motion detection thresholds for dichoptic plaids (Gorea et al., 2001; Maehara et al., 2017) has found evidence for monocular direction opponency, but not dichoptic opponency. Computational modeling of these results suggested that opponency occurs before divisive gain control operates (Maehara et al., 2017). Our results thus contrast with these findings in two ways - in the existence of dichoptic opponency and in how dichoptic sensitivity is being limited. The stimuli used in the psychophysical experiments were very different than ones used here, consisting of grating patches drifting at relatively high temporal frequencies (e.g. 7.5 or 20 Hz), that cannot support image segmentation on the basis of relative motion/disparity cues. Our stimuli can, and we found that opponent suppression depended strongly on the presence of a reference. At 5 months of age, we see no evidence of the dichoptic opponent process we observed for adults. These results, combined with previous results with CDOT-isolating stimuli (Norcia et al., 2017a), indicate that the binocular cues of IVOD and CDOT, each capable - in principle - of supporting MID, are both functionally immature.

### Implications for models of relative motion coding

Our results suggest several possible extensions to existing models of motion and disparity processing. Motion responses in V1 have traditionally been modeled with variants of an energy-like metric (Adelson and Bergen, 1985; van Santen and Sperling, 1985; Watson and Ahumada, 1985) that were not designed to explain human observers’ greatly enhanced sensitivity to relative motion (Legge and Campbell, 1981; Nakayama and Tyler, 1981; McKee et al., 1990). Motion energy models could be extended along the lines of an existing model of relative disparity processing that pools the outputs of analogous disparity energy detectors in V1 at a second stage (Bredfeldt et al., 2009). The model combines V1 disparity energy units in an opponent fashion and is successful in explaining several properties of V2 cell responses to disparity edges. Formulating a relative motion model along these lines, with motion energy substituted for disparity energy at the first stage, would certainly be feasible and thus the effects of references in both motion and disparity domains could be accommodated by an analogous two-stage model. Existing physiological and fMRI data suggest that both stages are present in V1 for motion (Reppas et al., 1997; Cao and Schiller, 2003; Kohler et al., 2017), but that the second stage of the relative disparity system begins in V2 or beyond (Peterhans and von der Heydt, 1993; Thomas et al., 2002; Bredfeldt and Cumming, 2006; Kohler et al., 2017). Prior data suggest that for both motion (the present results) and disparity (Norcia et al., 2017a) this hypothesized second stage is immature in infants. In the case of motion, non-classical surround effects (Bridgeman, 1972; Allman et al., 1985; Born, 2000; Cao and Schiller, 2003; Shen et al., 2007) are viable candidates for an underlying mechanism for enhanced responses to relative motion. Analogous non-classical surround effects in the disparity domain have been observed in macaque MT (Bradley and Andersen, 1998) and MST (Eifuku and Wurtz, 1999). Human fMRI data also provides evidence for opponency in the disparity domain, starting no later than V3 (Kohler et al., 2017).

### Implications for models of relative disparity coding

Disparity tuning in macaque V1 is distributed over all orientations and is thus not specifically associated with horizontal disparities that support stereopsis (Cumming, 2002; Read and Cumming, 2004). Relative disparity tuning in single cells, to our knowledge, has only been probed with horizontal disparities (Janssen et al., 2001; Thomas et al., 2002; Umeda et al., 2007; Anzai et al., 2011; Krug and Parker, 2011; Shiozaki et al., 2012). The present results suggest that sensitivity to relative *vertical* disparity is present at least for larger disparities presented over a relatively large field-of-view (up to 20 deg eccentricity). Our neural correlate of relative disparity processing is the first harmonic response. We showed that the first harmonic arises from CDOT rather than an IOVD because the response is eliminated in the two stimulus conditions (inter-ocularly uncorrelated and anti-correlated dots) that remove percepts of MID from CDOT, but not from IOVD. Functionally, our candidate vertical relative disparity signal is robust when a reference is present and is eliminated when the reference is removed, as is the case for horizontal disparity. Further corroboration comes from Experiment 3 where disparity modulates symmetrically around the reference. This manipulation specifically varies the relative disparity between the reference and moving bands as well as the global stimulus configuration. Uniform and segmented states could thus be differentiated based on the first harmonic signal for both horizontal and vertical stimulus orientations. A second-stage pooling of vertical absolute disparities could give rise to vertical relative disparity sensitivity in the same way as has been suggested for horizontal relative disparity (Bredfeldt et al., 2009). Computationally, this model implements a form of opponency in the disparity domain, but the opponency is within the classical receptive field of second stage V2 neurons, rather than between classical receptive field and its surround. Whether vertical relative disparity sensitivity is a property of the classical receptive field or center-surround interactions remains to be determined.

A characteristic behavior that any model of relative disparity processing will need to account for is the fact that sensitivity is dramatically greater for horizontal compared to vertical relative disparity by a factor of at least 4 in the adults (Figure 6). Whether the difference is due to the properties of first-order cells in V1 or whether it represents a specific adaptation to horizontal disparities in higher visual areas remains to be determined.

### Implications for models of MID encoding

Different models of MID have compared architectures where disparity is extracted first, followed by a second stage that computes changes in disparity over time (CDOT) with architectures where velocity is computed first and then differenced at a second stage (IOVD; Cumming and Parker, 1994; Peng and Shi, 2010, 2014). We have presented evidence for each of these processes. The existing models were conceived in the context of MID from horizontal disparity/motion. Our IOVD correlate, the suppression of the second harmonic response for anti-phase motion, is present for both horizontal and vertical displays. The same is true for our CDOT correlate in the first harmonic. Thus, neither of these visual responses are specific to horizontal motion or a percept of MID. This suggests that CDOT and IOVD, processed in isolation or combined, are insufficient to produce a percept of MID. Rather, we have identified what appears to be an intermediate set of signal processing operations that either occur before MID extraction or that operate in parallel with it. Moreover, none of the existing models of CDOT or IOVD take into account the role of references that we show strongly influence the strength of responses.

Which responses in our data are likely to reflect activity specifically related to MID? Evidence for a MID signal based on CDOT comes from our finding that disparity thresholds for the CDOT-specific response at the first harmonic is much lower for horizontal than for vertical conditions, consistent with the prominent role that horizontal relative disparity plays in perception. We see very little evidence for an asymmetry between horizontal and vertical conditions at the second harmonic, our IOVD correlate. We do, however see a measurable superiority of responsiveness to smaller horizontal displacements at the fourth harmonic that persists for stimuli that isolate the IOVD cue. The fact that the fourth harmonic - but not the second harmonic - is tuned for stimulus orientation indicates that the two harmonics are not being generated by a common, higher-order non-linearity, such as rectification in a population of transient or direction-selective cells. An alternative view that is qualitatively consistent with velocity-first MID models is a cascade model in which a first stage motion computation creates second harmonics at its output that are then pooled in a non-linear fashion by a binocular MID process, with the result being a response that is fourth order with respect to the input. In this model, the second harmonic would have to be monocular for the fourth harmonic to represent the output of the binocular MID stage. However, the second harmonic is modulated by inter-ocular phase and is thus at least partly due to binocular mechanisms. This suggests that MID and anti-phase suppression may arise in parallel pathways, rather than being properties of a common MID mechanism.

### Implications for models of human visual development

The visual evoked responses of 5-month-old infants in the presence of high-quality motion and disparity references bear a strong resemblance to the unreferenced responses of adults, suggesting that both relative motion and relative disparity mechanisms are selectively immature in infants. Relative motion sensitivity is quantitatively immature, being ∼8 times lower than that of the adult (compare Figure 4 to Figure 3A). The present results showing immature relative disparity mechanisms at this age are consistent with our previous results from CDOT-isolating dynamic random dot stereograms (Norcia et al., 2017a).

In addition to their lack of a sensitivity to references, infants also show a lack of anti-phase suppression. Because anti-phase suppression in adults is strongly dependent on the presence of references, it is conceivable that the lack of anti-phase suppression in infants is driven by their lack of sensitivity to references. However, it is also possible that the lack of anti-phase suppression represents a separate immaturity relating to binocular vision. In-phase motion responses were measured under binocular conditions, but would likely be very similar if presented under monocular viewing conditions. Thus, sensitivity to references is not inherently a binocular phenomenon, whereas anti-phase suppression is. The infant visual system thus displays qualitative immaturities and not just quantitative differences in its sensitivity to displacement.

## Materials and Methods

### Participants and Procedure

A total of 59 adults participated in one or more of the experiments (age range 17.1 to 40.8 years; mean 23.2, SD = 5.24). Twenty-two healthy full-term infants with birthweights exceeding 2500g participated (12 male, avg. age = 5.6 months, SD = 1.1). The adults had normal or corrected-to-normal vision, with a visual acuity of 0.1 LogMar in each eye or better and no significant difference of performance between both eyes. Adult participants also scored at least 40 arcsec or better on the RanDot Stereogram test. Prior to the experiment, the procedure was explained to each participant or the parent, and written informed consent was obtained before the experiment began. The protocol was approved by the Stanford University Institutional Review Board. Adult participants were solicited though Stanford University subject pools. Infants were recruited via mailers sent to addresses procured by the California Department of Public Health, Center for Health Statistics and Informatics.

### Stimuli

In all experiments, participants viewed Random Dot Kinematograms (RDK) or stereograms (RDS) on a 65-inch Sony Bravia XBR-65HX929 LCD monitor. The dots were drawn with OpenGL using antialiasing at a screen resolution of 1920 × 1080 pixels. This function allowed us to present disparities via dithering that were smaller than the nominal resolution set by the 1920 × 1080 display matrix. This was verified by examining the contents of video memory and through examination of the anti-aliased pixels under magnification. The dots were updated at 20 Hz.

For 7 of 8 experiments, viewing distance was set at 1 m, resulting in a 40 × 40 deg field of view, a 4.5 arcmin dot diameter, with 5 dots per square degree. In the first psychophysics experiment, the viewing distance was set at 3 m, to allow for smaller increments in dot locations. The stimuli were rendered as Red/Blue anaglyphs. The luminance of the images in the two eyes was equated by calibrating the display through each filter. A −0.5 diopter lens was placed in the Blue channel of the adult glasses to compensate for the differential focus caused by chromatic aberration. Cross-talk was minimal perceptually when viewing high-contrast images in the two eyes.

### Experimental design and procedure

Schematic illustrations of the stimuli are provided in Figure 1. The displays in each Experiment comprised a set of alternating bands containing random dots that could differ in their inter-ocular correlation, temporal coherence or both. One set of “test bands” underwent in-phase or anti-phase motion on each trial, with the adjacent “reference bands” serving as a static or dynamic reference. There were 20 test bands and 20 reference bands on the screen (spatial frequency of ∼0.45 c/deg). When the inter-ocular correlation was 100%, there were matching dots in each eye, when it was 0%, independently generated dots were presented to each eye. When the inter-ocular correlation was −100%, the dark dots in one eye were matched with bright dots in the other eye (see Experiment 5 for further details). Displays in which the temporal correlation was 100% had dots that moved coherently with unlimited lifetime. Displays that had a temporal coherence of 0% had newly generated dots on every image update (20 Hz). Dots were replotted at the end of motion trajectory to keep the number of dots on the screen the same at all times and the location of the borders of the dot region constant.

Experiments 1, 2, and 3 used a 2 × 2 × 2 design, with factors of inter-ocular phase, reference quality and stimulus orientation, resulting in 8 conditions. Inter-ocular phase of the moving test bands was either in-phase or in anti-phase, *e.g*. either a 0 or 180 deg temporal phase relationship applied between eyes. Manipulation of the dots in the reference region created referenced and unreferenced motion and disparity conditions. In Experiments 4 and 5, manipulation of the inter-ocular correlation in the moving bands (changing them from 100% correlated between eyes to either 0% or −100%) was used to isolate the IOVD cue. Here the design was also 2 × 2 × 2 with factors of inter-ocular phase, inter-ocular correlation and orientation. In each of the adult experiments, participants completed 15 trials per stimulus condition, and the 8 conditions were run in a block-randomized fashion: 5 consecutive trials of a given condition comprised a block. Data were collected in 3 separate, continuous recording sessions, each lasting approximately 8 minutes, with a break given in between. During each session, a single block of each condition was run. Observers were instructed to fixate the center of the display and to withhold blinks. For the infant experiment, only the four horizontal conditions were used, resulting in a 2 × 2 design, with factors of inter-ocular phase and reference quality. Infants completed 10 trials. Details of each experiment are provided below.

### Experiment 1: Near vs Zero, absolute motion with noise reference

Twelve adults (5 male) with an average age of 22.2 years (SD = 4.97) participated. Participants viewed horizontal and vertical stimulus orientations at 1 meter that were either in-phase (2D motion) or anti-phase. For the horizontal orientation, the anti-phase condition created crossed disparities in the test bands that alternated with zero disparity at 2 Hz. Zero disparity was at the plane of the display. Stimulus displacement in each eye swept from 0.5 to 16 arcmin in 10 equal log steps, which were updated at 1 second intervals. Trials lasted 12 seconds, with the first and last step being repeated at the beginning and end of each trial, respectively. This was done to minimize effects of contrast transients when the dots first appeared, and to ensure that participants did not blink during the middle 10 sec of each trial, which went into the data analysis. All displacements are plotted as the single-eye displacement value. The displacement was modulated with a square wave temporal profile. The reference bands were static in the relative motion conditions, and contained incoherent, uncorrelated motion in the absolute motion conditions.

### Experiment 2: Near vs Zero, absolute motion with blank reference

Thirteen adults (6 male, avg. age = 22.9 years, SD = 4.92) participated. Blank reference bands (dark empty regions with no dots) replaced incoherent motion reference bands for the absolute motion conditions. All other stimulus features from Experiment 1 were kept constant.

### Experiment 3: Near vs Far, absolute motion with noise reference

Twelve adults (7 male, avg. age=22.1 years, SD=5.08) participated. The end-points of the motion trajectories were set so that for the anti-phase horizontal displacement conditions, the disparity alternated between equal crossed and uncrossed values about zero disparity. The magnitude of peak to peak-disparity in the test bands matched that of all other experiments, only the disparity of the test bands relative to that of the reference bands was changed. Absolute motion conditions presented incoherent motion in the reference bands, as in Experiment 1.

### Experiment 4: Uncorrelated test: IOVD isolation

Thirteen adults (6 male, avg. age = 24.6 years, SD = 5.14) participated. The relative motion and relative disparity conditions were the same as in Experiment 1, but the absolute motion conditions were replaced with conditions in which moving test bands contained uncorrelated dots while reference bands contained static dots.

### Experiment 5: Anti-correlated test: IOVD isolation

Thirteen adults (7 male, avg. age = 27.3 years, SD = 6.9) participated. The relative motion and relative disparity conditions were similar to those used in Experiment 1, but the absolute motion conditions were replaced with conditions in which moving test bands contained anti-correlated dots while reference bands contained static dots. In all conditions, dots were presented on a mean-luminance purple background, which allowed bright and dark dots for both red and blue color channels to be shown to either eye in the anti-correlated display.

### Psychophysics Experiment 1 with relative displacement conditions

Fourteen adults (7 male, avg. age = 25.8 years, SD = 5.87) participated. Viewing distance was set at 3m rather than the 1m used for all other experiments to allow for smaller displacement increments. Spatial frequency of the static and reference bands was adjusted to 0.46 cpd and dot diameter to 4.2 arcmin to match these parameters with those used at 1m. Displacements ranged from 0.16 to 2.56 arcmin in 10 equal log steps. Parameters were otherwise matched to the relative conditions of Experiments 1 and 2, with static reference bands. On each trial, participants viewed either an ascending or descending sweep for a given condition and were instructed to press the right arrow key on a keyboard button whenever they detected a state change in the stimuli. For ascending sweeps, participants pressed the key when they first perceived the dots to change from static to moving. For descending sweeps, participants pressed the key when they perceived the moving dots changed to static. Data from 2 participants were excluded from analysis because they failed to respond or responded incorrectly in more than 15% of trials.

### Psychophysics Experiment 2 with absolute conditions

Eleven adults (6 male, avg. age = 24.8 years, SD = 5.9) participated. 5 participants (3 male, avg. age = 29.1, SD = 4.6) took part in both psychophysics experiments. The display parameters matched the absolute conditions of Experiment 2, with blank reference bands, except that participants were shown both ascending and descending sweeps. Participants completed the same behavioral measure as described for Psychophysics Experiment 1. We also included infrequent catch trials for ascending and descending conditions, in which the state of motion never changed, i.e. the dots remained static or moving for the duration of the trial. Participants were shown these types of catch trials and understood there should be no response when encountered during the recording session. Catch trials were excluded from the analysis. Data from 1 participant were excluded from analysis because they failed to respond or responded incorrectly in more than 15% of trials.

### Infant Experiment

Infants completed a two-part, reduced protocol that consisted of the four horizontal conditions from Experiment 2, but with a displacement range of 2-32 arcmin. The adjusted sweep was used to compensate for the infants’ elevated displacement threshold (Norcia et al., 2017b). Infants viewed the horizontal relative motion conditions (in-phase and anti-phase) and blank reference absolute motion conditions (in-phase and anti-phase) during separate visits. Condition order was counterbalanced for session one and two. Out of 22 infants, 4 were excluded from the analysis because they were unable to complete at least 5 trials of each condition. Sixteen infants (7 male, avg. age = 5.3 months, SD = 0.84) completed both the relative motion and absolute motion session, while another 2 completed only the absolute motion session, for a total of 18 infant datasets (8 male, avg. age =5.32 months, SD = 0.80).

#### EEG Acquisition and Processing

Data were collected from all participants using high-density HydroCell electrode arrays paired with an Electrical Geodesics NetAmp400 and accompanying NetStation 5 software. The nets used for adults had 128 channel and the one used for infants had 124. EEG data initially sampled at 500 Hz was resampled at 420 Hz to provide 7 samples per video frame. Digital triggers were sent from in-house stimulus presentation software and stored with the EEG recording to allow synching of the visual stimulus and EEG with millisecond precision. Recordings were exported from NetStation using a 0.3-50 Hz bandpass filter, which was applied twice to ensure that power in frequencies outside the filter range was eliminated. The data were then imported into in-house signal processing software for preprocessing. If more than 15% of samples for a given sensor exceeded an amplitude threshold, the sensor was excluded from further analysis. Adult data were evaluated using a 30 μV threshold, whereas a more liberal threshold was applied for the infant data, ranging between 30 and 100 μV. Sensors that were noisier than the threshold were replaced by an averaged value from six of their nearest neighbors. The EEG data was then re-referenced to the common average of all the sensors and segmented into ten 1000 ms long epochs (each corresponding to exactly 2 stimulus cycles). Epochs for which more than 10% of data samples exceeded a noise threshold (30 μV for both adult and infant participants) or for which any sample exceeded a peak/blink threshold (60 μV for both adult and infant participants) were excluded from the analysis on a sensor-by-sensor basis. If more than 7 sensors had epochs that exceeded the peak/blink threshold, the entire epoch was rejected for all channels. This was typically the case for epochs containing artifacts, such as blinks or eye movements.

#### Data Analysis

In our sweep paradigm, stimulus values were updated for every 1-sec bin, so each epoch in our analysis is tied to a distinct set of stimulus parameters, for a given trial. The amplitude and phase of the Steady-State Visual Evoked Potentials (SSVEPs) were extracted using a Recursive Least Squares adaptive filter (Tang and Norcia, 1995) with a memory length equal to the 1 sec bin length. Real and imaginary components of the SSVEPs at the first four harmonics of the stimulus frequency were calculated. Noise estimates were calculated at neighboring frequency bins, i.e. F +/- 1 Hz.

We reduced the spatial dimensionality of our data by decomposing the sensor data into a set of physiologically interpretable components using Reliable Components Analysis (RCA; Dmochowski et al., 2015). Because SSVEP response phase is constant over repeated trials of the same stimulus, RCA utilizes a cross-trial covariance matrix to decompose the 128-channel montage into a smaller number of components that maximize trial-to-trial consistency through solving for a generalized eigenvalue problem. The real and imaginary values for each 1 sec epoch, across the 128 sensors, and across trials and participants, served as the input data for RCA. Reliable components were derived separately for each harmonic in each of the five main experiments, and separately for each harmonic in the infant experiment. The data reduction for the SSVEP data collected during the psychophysical experiments was done by projecting the data through the component weights from the first RC derived for *2F* in Experiment 2, which used the same stimulus parameters as the psychophysical experiments.

We ran two additional reliable components analyses combining data from unique participants across several experiments. One exclusively with data from the static reference horizontal and vertical anti-phase conditions from Experiments 1, 2, 3 and 4, yielding a data set with 39 unique participants, and another with data from the horizontal and vertical anti-phase conditions in the uncorrelated and anti-correlated IOVD-isolating Experiments 4 and 5, yielding a data set with 22 unique participants. In both cases, we derived reliable components separately for the *2F* and *4F* data, using the same RCA approach as for the individual experiments.

Our analyses focused on the first RC component, which for *2F* data explained much of the reliability in the 5 main experiments (average = 67.8%, SD = 3.1) and a substantial amount of the variance (average = 23.1% SD = 1.2). This was also the case for *1F* for 4 out of 5 experiments, excluding Experiment 3 where *1F* were not measurable (average reliability explained average = 49.3% SD = 3.9; average variance explained = 18.6% SD = 3.5). For the infant *1F* data, the first RC did not look like a visual response, likely because SSVEPs were weak overall. We did however see a topography and response function that resembled that observed for adults in RC5, which we present in Figure 7.

After projecting the epoch-level data through the RCA component weights, averages across trials in each condition and across participants were computed by averaging the real and imaginary coefficients for a given response harmonic (vector average responses). The averages were computed separately for each of the 10 bins in the displacement sweeps. Noise estimates based on neighboring frequency bins did not contribute to RCA but were projected through the component weights to allow comparison with the SSVEP data, and then averaged across trials, participants and conditions.

We fit the vector-averaged response functions with the following equation (Naka-Rushton function) (Naka and Rushton, 1966).

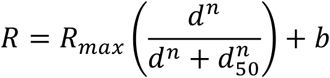

Where *R* is the response, *d* is the displacement of the moving bands, and *b* is the baseline. *R_max_* (maximal response), *n* (exponent of the power function, > 0), *b* and *d_50_* (displacement at half *R_max_)* are free parameters. We computed standard errors for each parameter based on a jackknife procedure in which the function was fitted to average data from all participants except one (Equation 2; Miller et al., 2009). We tested whether the fit parameters were significantly different across conditions by computing *t*-values based on the jackknifed standard error of the difference (Equations 2 and 3; Miller et al., 1998).

## Acknowledgements

This research was supported by National Institutes of Health grant EY018775.

## References

Adelson EH, Bergen JR (1985) Spatiotemporal energy models for the perception of motion.J Opt Soc Am A 2:284–299.

Allman J, Miezin F, McGuinness E (1985) Direction- and velocity-specific responses from beyond the classical receptive field in the middle temporal visual area (MT). Perception 14:105–126.

Anzai A, Chowdhury SA, DeAngelis GC (2011) Coding of Stereoscopic Depth Information in Visual Areas V3 and V3A. The Journal of Neuroscience 31:10270–10282.

Bakin JS, Nakayama K, Gilbert CD (2000) Visual responses in monkey areas V1 and V2 to three-dimensional surface configurations. The Journal of neuroscience: the official journal of the Society for Neuroscience 20:8188–8198.

Beverley KI, Regan D (1973) Evidence for the existence of neural mechanisms selectively sensitive to the direction of movement in space. The Journal of physiology 235:17–29.

Born RT (2000) Center-surround interactions in the middle temporal visual area of the owl monkey. J Neurophysiol 84:2658–2669.

Bradley DC, Andersen RA (1998) Center-surround antagonism based on disparity in primate area MT. J Neurosci 18:7552–7565.

Bredfeldt CE, Cumming BG (2006) A simple account of cyclopean edge responses in macaque v2. J Neurosci 26:7581–7596.

Bredfeldt CE, Read JC, Cumming BG (2009) A quantitative explanation of responses to disparity-defined edges in macaque V2. J Neurophysiol 101:701–713.

Bridgeman B (1972) Visual receptive fields sensitive to absolute and relative motion during tracking. Science (New York, NY 178:1106–1108.

Brooks KR, Stone LS (2004) Stereomotion speed perception: Contributions from both changing disparity and interocular velocity difference over a range of relative disparities. Journal of vision 4:6–6.

Brooks KR, Stone LS (2006) Stereomotion suppression and the perception of speed: accuracy and precision as a function of 3D trajectory. Journal of vision [electronic resource] 6:1214–1223.

Cao A, Schiller PH (2003) Neural responses to relative speed in the primary visual cortex of rhesus monkey. Visual neuroscience 20:77–84.

Carandini M (2012) From circuits to behavior: a bridge too far? Nature neuroscience 15:507–509.

Cooper EA, van Ginkel M, Rokers B (2016) Sensitivity and bias in the discrimination of two-dimensional and three-dimensional motion direction. Journal of vision [electronic resource] 16:5.

Cormack LK, Czuba TB, Knoll J, Huk AC (2017) Binocular Mechanisms of 3D Motion Processing. Annu Rev Vis Sci 3:297–318.

Cottereau BR, McKee SP, Norcia AM (2014) Dynamics and cortical distribution of neural responses to 2D and 3D motion in human. J Neurophysiol 111:533–543.

Cumming BG (2002) An unexpected specialization for horizontal disparity in primate primary visual cortex. Nature 418:633–636.

Cumming BG, Parker AJ (1994) Binocular mechanisms for detecting motion-in-depth. Vision research 34:483–495.

Cumming BG, Parker AJ (1997) Responses of primary visual cortical neurons to binocular disparity without depth perception. Nature 389:280–283.

Cumming BG, Parker AJ (1999) Binocular neurons in V1 of awake monkeys are selective for absolute, not relative, disparity. J Neurosci 19:5602–5618.

Cumming BG, Parker AJ (2000) Local disparity not perceived depth is signaled by binocular neurons in cortical area V1 of the Macaque. J Neurosci 20:4758–4767.

Czuba TB, Rokers B, Huk AC, Cormack LK (2010) Speed and eccentricity tuning reveal a central role for the velocity-based cue to 3D visual motion. J Neurophysiol 104:2886–2899.

Dmochowski JP, Greaves AS, Norcia AM (2015) Maximally reliable spatial filtering of steady state visual evoked potentials. NeuroImage 109:63–72.

Dow BM (1974) Functional classes of cells and their laminar distribution in monkey visual cortex. J Neurophysiol 37:927–946.

Dubner R, Zeki SM (1971) Response properties and receptive fields of cells in an anatomically defined region of the superior temporal sulcus in the monkey. Brain research 35:528–532.

Eifuku S, Wurtz RH (1999) Response to motion in extrastriate area MSTl: disparity sensitivity. J Neurophysiol 82:2462–2475.

Frost BJ, Nakayama K (1983) Single visual neurons code opposing motion independent of direction. Science (New York, NY 220:744–745.

Glennerster A, McKee SP (1999) Bias and sensitivity of stereo judgements in the presence of a slanted reference plane. Vision research 39:3057–3069.

Glennerster A, McKee SP, Birch MD (2002) Evidence for surface-based processing of binocular disparity. Curr Biol 12:825–828.

Gorea A, Conway TE, Blake R (2001) Interocular interactions reveal the opponent structure of motion mechanisms. Vision research 41:441–448.

Harris JM, Nefs HT, Grafton CE (2008) Binocular vision and motion-in-depth. Spatial vision 21:531–547.

Hubel DH, Wiesel TN (1968) Receptive fields and functional architecture of monkey striate cortex. The Journal of physiology 195:215–243.

Janssen P, Vogels R, Liu Y, Orban GA (2001) Macaque inferior temporal neurons are selective for three-dimensional boundaries and surfaces. J Neurosci 21:9419–9429.

Katz LN, Hennig JA, Cormack LK, Huk AC (2015) A Distinct Mechanism of Temporal Integration for Motion through Depth. The Journal of Neuroscience 35:10212–10216.

Kohler PJ, Cottereau B, Norcia AM (2017) Image segmentation based on relative motion and relative disparity cues in topographically organized areas of human visual cortex. bioRxiv.

Krug K, Parker AJ (2011) Neurons in dorsal visual area V5/MT signal relative disparity. J Neurosci 31:17892–17904.

Lages M, Heron S (2010) On the inverse problem of binocular 3D motion perception. PLoS Comput Biol 6:e1000999.

Legge GE, Campbell FW (1981) Displacement detection in human vision. Vision research 21:205–213.

Maeda M, Sato M, Ohmura T, Miyazaki Y, Wang AH, Awaya S (1999) Binocular depth-from-motion in infantile and late-onset esotropia patients with poor stereopsis. Investigative ophthalmology & visual science 40:3031–3036.

Maehara G, Hess RF, Georgeson MA (2017) Direction discrimination thresholds in binocular, monocular, and dichoptic viewing: Motion opponency and contrast gain control. Journal of vision [electronic resource] 17:7.

Maunsell JH, Van Essen DC (1983) Functional properties of neurons in middle temporal visual area of the macaque monkey. I. Selectivity for stimulus direction, speed, and orientation. J Neurophysiol 49:1127–1147.

McKee SP, Welch L, Taylor DG, Bowne SF (1990) Finding the common bond: stereoacuity and the other hyperacuities. Vision research 30:879–891.

Mikami A, Newsome WT, Wurtz RH (1986) Motion selectivity in macaque visual cortex. II. Spatiotemporal range of directional interactions in MT and V1. J Neurophysiol 55:1328–1339.

Miller J, Patterson T, Ulrich R (1998) Jackknife-based method for measuring LRP onset latency differences. Psychophysiology 35:99–115.

Miller J, Ulrich R, Schwarz W (2009) Why jackknifing yields good latency estimates. Psychophysiology 46:300–312.

Naka KI, Rushton WAH (1966) S-potentials from luminosity units in the retina of fish (Cyprinidae). The Journal of Physiology 185:587–599.

Nakayama K, Tyler CW (1981) Psychophysical isolation of movement sensitivity by removal of familiar position cues. Vision research 21:427–433.

Nefs HT, Harris JM (2010) What visual information is used for stereoscopic depth displacement discrimination? Perception 39:727–744.

Norcia AM, Gerhard HE, Meredith WJ (2017a) Development of Relative Disparity Sensitivity in Human Visual Cortex. J Neurosci 37:5608–5619.

Norcia AM, Pei F, Kohler PJ (2017b) Evidence for long-range spatiotemporal interactions in infant and adult visual cortex. Journal of vision 17:12–12.

Orban GA, Kennedy H, Bullier J (1986) Velocity sensitivity and direction selectivity of neurons in areas V1 and V2 of the monkey: influence of eccentricity. J Neurophysiol 56:462–480.

Peng Q, Shi BE (2010) The changing disparity energy model. Vision research 50:181–192.

Peng Q, Shi BE (2014) Neural population models for perception of motion in depth. Vision research 101:11–31.

Peterhans E, von der Heydt R (1993) Functional organization of area V2 in the alert macaque. The European journal of neuroscience 5:509–524.

Petrov Y, Glennerster A (2004) The role of a local reference in stereoscopic detection of depth relief. Vision research 44:367–376.

Petrov Y, Glennerster A (2006) Disparity with respect to a local reference plane as a dominant cue for stereoscopic depth relief. Vision research 46:4321–4332.

Poggio GF, Gonzalez F, Krause F (1988) Stereoscopic mechanisms in monkey visual cortex: binocular correlation and disparity selectivity. J Neurosci 8:4531–4550.

Poggio GF, Motter BC, Squatrito S, Trotter Y (1985) Responses of neurons in visual cortex (V1 and V2) of the alert macaque to dynamic random-dot stereograms. Vision research 25:397–406.

Read JC, Cumming BG (2004) Understanding the cortical specialization for horizontal disparity. Neural Comput 16:1983–2020.

Reppas JB, Niyogi S, Dale AM, Sereno MI, Tootell RB (1997) Representation of motion boundaries in retinotopic human visual cortical areas. Nature 388:175–179.

Rokers B, Cormack LK, Huk AC (2008) Strong percepts of motion through depth without strong percepts of position in depth. Journal of vision [electronic resource] 8:6 1–10.

Rokers B, Cormack LK, Huk AC (2009) Disparity- and velocity-based signals for three-dimensional motion perception in human MT+. Nature neuroscience 12:1050–1055.

Samonds JM, Tyler CW, Lee TS (2017) Evidence of Stereoscopic Surface Disambiguation in the Responses of V1 Neurons. Cereb Cortex 27:2260–2275.

Schiller PH, Finlay BL, Volman SF (1976) Quantitative studies of single-cell properties in monkey striate cortex. I. Spatiotemporal organization of receptive fields. J Neurophysiol 39:1288–1319.

Serrano-Pedraza I, Phillipson GP, Read JC (2010) A specialization for vertical disparity discontinuities. Journal of vision [electronic resource] 10:2 1–25.

Shen ZM, Xu WF, Li CY (2007) Cue-invariant detection of centre-surround discontinuity by V1 neurons in awake macaque monkey. The Journal of physiology 583:581–592.

Shioiri S, Saisho H, Yaguchi H (2000) Motion in depth based on inter-ocular velocity differences. Vision research 40:2565–2572.

Shiozaki HM, Tanabe S, Doi T, Fujita I (2012) Neural Activity in Cortical Area V4 Underlies Fine Disparity Discrimination. The Journal of Neuroscience 32:3830–3841.

Tang Y, Norcia AM (1995) An adaptive filter for steady-state evoked responses. Electroencephalography and clinical neurophysiology 96:268–277.

Thomas OM, Cumming BG, Parker AJ (2002) A specialization for relative disparity in V2. Nature neuroscience 5:472–478.

Tyler CW (1971) Stereoscopic depth movement: two eyes less sensitive than one. Science (New York, NY 174:958–961.

Umeda K, Tanabe S, Fujita I (2007) Representation of Stereoscopic Depth Based on Relative Disparity in Macaque Area V4. Journal of neurophysiology 98:241–252.

van Santen JP, Sperling G (1985) Elaborated Reichardt detectors. J Opt Soc Am A 2:300–321.

von Grunau M, Frost BJ (1983) Double-opponent-process mechanism underlying RF-structure of directionally specific cells of cat lateral suprasylvian visual area. Experimental brain research Experimentelle Hirnforschung 49:84–92.

Watson AB, Ahumada AJ, Jr. (1985) Model of human visual-motion sensing. J Opt Soc Am A 2:322–341.

Westheimer G, McKee SP (1979) What prior uniocular processing is necessary for stereopsis? Investigative ophthalmology & visual science 18:614–621.

